# MACI: A machine learning-based approach to identify drug classes of antibiotic resistance genes from metagenomic data

**DOI:** 10.1101/2023.04.18.537418

**Authors:** Rohit Roy Chowdhury, Jesmita Dhar, Stephy Mol Robinson, Abhishake Lahiri, Sandip Paul, Kausik Basak, Rachana Banerjee

**Author notes:** Corresponding Author: Dr. Rachana Banerjee.

## Abstract

Novel methodologies are now essential for identification of antibiotic resistant pathogens in order to resist them. Here, we are presenting a model, MACI (Machine learning-based Antibiotic resistance gene-specific drug Class Identification) that can take metagenomic fragments as input and predict the drug class of antibiotic resistant genes. We trained the model to learn underlying patterns of genes using the Comprehensive Antibiotic Resistance Database. It comprises of 116 drug classes with a total of 2960 representative sequences. Among these 116 drug classes, we found 22 categories (contributing approximately 85% of the overall sequence-data) surpassed other 94 drug classes based on the number of fragments. The model showed an average precision of 0.83 and a recall of 0.81 for these 22 drug classes. Moreover, the model predicted multidrug resistant classes with higher performance score (precision and recall: 0.9 and 0.88 respectively) compared to single drug resistant categories (0.77 and 0.75). Post to this, we analysed these 22 drug classes to find out class-specific overlapping patterns of nucleotides that led to accurate classification. This way, we found five drug classes viz. “carbapenem;cephalosporin;penam”, “cephalosporin”, “cephamycin”, “cephalosporin;monobactam;penam;penem”, and “fluoroquinolone”. Additionally, the positions of these significant patterns corroborated with the functional domains of majority of antibiotic resistance genes in that drug class, indicating their biological importance. These class-specific patterns play a pivotal role in rapid identification of some drug classes comprising antibiotic resistance genes. Further analysis showed that bacterial species, containing these five-drug classes, were found to have well-known multidrug resistance property.

## 1. Introduction

Over a past few years, researches in bioinformatics have thrusted an eye-awakening concern towards antibiotic resistance due to its significant impact in global public health. According to various studies, the fatal effect accounted for antibiotic resistance was estimated to be around 0.7 million each year globally. The same was expected to grow to a figure of 10 million by 2050, if timely measures are not taken [1–3]. In countries where standard treatment procedures are not properly followed, it is accelerating due to misuse and overuse of various antibiotics. This has led to a vulnerable situation in which the treatment of common infectious diseases is facing a great challenge. The growing cases of infections like pneumonia, tuberculosis, gonorrhoea, foodborne diseases etc. are strictly pointing to the resistance of various bacterial species with respect to a plethora of antibiotics. In 2015, World Health Organization (WHO) commenced an initiative by devising a global action plan to decipher the emergence of new resistance mechanisms. Some primary care strategies to effectively prevent the spreading of infectious diseases were also proposed [https://www.who.int/news-room/fact-sheets/detail/antibiotic-resistance]. Classically, antibiotic resistance can be acquired either by horizontal transfer of genes or by the mutations in preceding DNA. Presently, there is a huge assortment of antibiotics present in the environment, to which microbes are constantly exposed. Co-existence of them leads to the generation of resistance genes in those microbes by horizontal gene transfer. Finally, when such resistant bacteria make way into animals and plants, either directly or via commensal bacteria, the antibiotic resistance genes (ARGs) are transferred to the former [1-6]. For example - methicillin-resistant *Staphylococcus aureus* strains accompany several infections and it was reported to show extreme drug-resistance [7]. *Mycobacterium tuberculosis* strains are multidrug resistant due to their inhibiting property towards rifampicin, fluoroquinolone and isoniazid [8]. Moreover, *Escherichia coli* has become colistin-carbapenem-resistant after acquiring ARGs like mcr-1 and blaNDM-1 [9, 10]. Since, the emergence of such antimicrobial resistant organisms becomes habitual in clinical settings, fast and accurate assessment of antibiotic susceptibility carries significant importance to narrow-down patient-centric administration of antibiotics.

Various researchers spotted ample amounts of ARGs in several environments like sludge, soil, wastewater, human gut, animal waste etc., signifying these atmospheres to be the reservoirs of ARGs [11, 12]. High loads of ARGs, in an explicit environment, indicate the prevalence of antibiotic resistant bacterial strains [13]. The existence of antibiotic resistome in these environments is a cause of global public health hazard. Therefore, exhaustive investigations of the richness and diversity of bacterial communities, along with their ARG reservoirs, are fundamental to portray a complete depiction of the potential threat and crucial for monitoring antibiotic resistance. Shotgun metagenomic techniques are frequently used to measure ARGs in the environment as it can produce information on entire gene pool of the bacterial communities. Moreover, metagenomic technique has the unique advantage of detecting rare ARGs and is independent of cultivability of the individual microbes [14, 15]. With the advent of high-throughput sequencing technologies, the study of ARGs in metagenomic samples has become much more rapid and sensitive. Numerous bioinformatic pipelines were devised in this context. Most of these pipelines, at first, perform whole genome assembly by aligning raw metagenomic reads with respect to accessible online databases containing whole genome sequences. Then, gene identification and annotation are carried out using some traditional methods like BLAST [16], Bowtie [17], or DIAMOND [18]. Such homology search-based methods use sequence similarity cut-off, based on which various categories of ARGs can be predicted or assigned. ARDB [19], SARG [20], CARD [21], and ResFinder [22] are a few of the reference databases which were constructed for homology-search. However, these databases contain only a small share of the total resistome. Also, in many cases, these databases are constructed under a specific genomic context which limits the search for ARG sequences in other contexts. As an example, for envisioning plasmid-borne ARGs, bioinformatics pipelines like ResFinder [22] and SEAR

[23] are used as the reference databases. Similarly, only 12 types of antimicrobials are identified by the Mykrobe predictor [24]. PATRIC can only identify the carbapenem, methicillin, and beta lactam resistant ARGs [25]. Alternatively, “best-hit” based methods are highly effective for detection of recognized and highly conserved ARGs. However, they suffer from the lack of a globally accepted, standardised similarity cut-offs. Too stringent cut-off leads to many false negatives, whereas, too lenient cut-off produces many false positives. Even without the cut-off issue, these methods suffer from several other disadvantages. Firstly, this pairwise alignment-based method is insensitive to point mutations. Subsequently, it may overlook novel genes. Secondly, the users have to optimize the other parameters used by the program very efficiently for correct prediction. Finally, the alignment-based methods rely on some curated database, which limits their precision to find out novel ARGs, especially from metagenomic reads [27]. Additionally, the prediction of whole genome from metagenomic reads (at the first step) can be highly time consuming and needs huge computational storage facilities.

The limitations, stemming from the existing bioinformatics methodologies, can be addressed using machine learning-based approaches. Machine learning-based computational models can eventually learn the underlying characteristics of ARGs from curated databases and make predictions about novel ARGs directly from metagenomic reads. In recent era, incorporation of complex mathematical reasoning has made the framework a powerful tool which imparts learning of significant patterns at multiple network levels. Different machine learning-based method viz. DeepARG, fARGene, HMD-ARG, GROOT were developed which can recognize ARGs directly from metagenomic short reads [28–30]. The computational framework facilitated more accurate identification of formerly uncharacterized resistance genes. Therefore, even if the unknown ARGs (having low sequence similarity to known ARGs) cannot be sensed using similarity-based methods, they can be fruitfully categorized using these machine learning-based models. However, the biological relevance associated with the identified sequence patterns were not established in these works [31, 32].

In this study, we intend to overcome these challenges through construction of a novel machine learning-based model for ARG-specific drug class identification (MACI). In a broader scene, our proposed model is an artificial neural network (ANN)-based algorithm which is deep in architecture. At different stages of the hidden layer network, it can capture significant patterns that lead to accurate classification of drug categories. It can take fragments of metagenomic sequences as input, train the network based on the diversity of genomic sequences, and predict the drug classes. Post to the curation of ARG sequence data, the whole sequence reads were fragmented into short length, encoded, and fed to the computational framework for training and optimization of network parameters. Once properly trained, the model was tested on remaining testing data to measure its efficacy in identifying the drug classes. These results were validated against the standard tools to justify the potentiality of the developed model. Further, in this work, we also established the biological significance of class-specific overlapping nucleotide patterns. MACI circumvents the computational needs for alignment based brute force matching or genome assembly steps.

## 2. Materials and Methods

Schematic of the developed model, MACI, is depicted in Fig. 1. For this work, we utilized the well-known Comprehensive Antibiotic Resistance Database (CARD) for ARG sequence retrieval [33, 34]. Recently, a comparative analysis of 12 ARG databases was performed and it was found that CARD covers high number of genes [35]. Therefore, it can be treated as a well-established resource of sequence data. The Resistance Gene Identifier (RGI) is the annotation tool which is used by CARD. This tool is available both as web-interface and downloadable command-line software. The ARG sequences (acquired from CARD) were first checked for redundancy removal to identify potentially discriminating sequences corresponding to different drug classes. Post to fragmentation of the sequences and data pre-processing, the dataset was divided into training (80%) and testing (20%) groups. An ANN-based supervised learner was developed and trained using the training data. The testing counterpart was used for performance measurement of the model. Once properly trained and tested, the computational framework facilitated accurate characterization of the drug classes of ARGs. In this work, we also identified class-specific overlapping patterns that led to accurate classification of drug classes. This was further explored to perform a correlation study with the taxonomic clades of ARGs. It enabled the understanding of biological significance of drug class-specific taxonomic variations. Step-wise detailed description of the methodology is given in the following part.

**Fig. 1.**
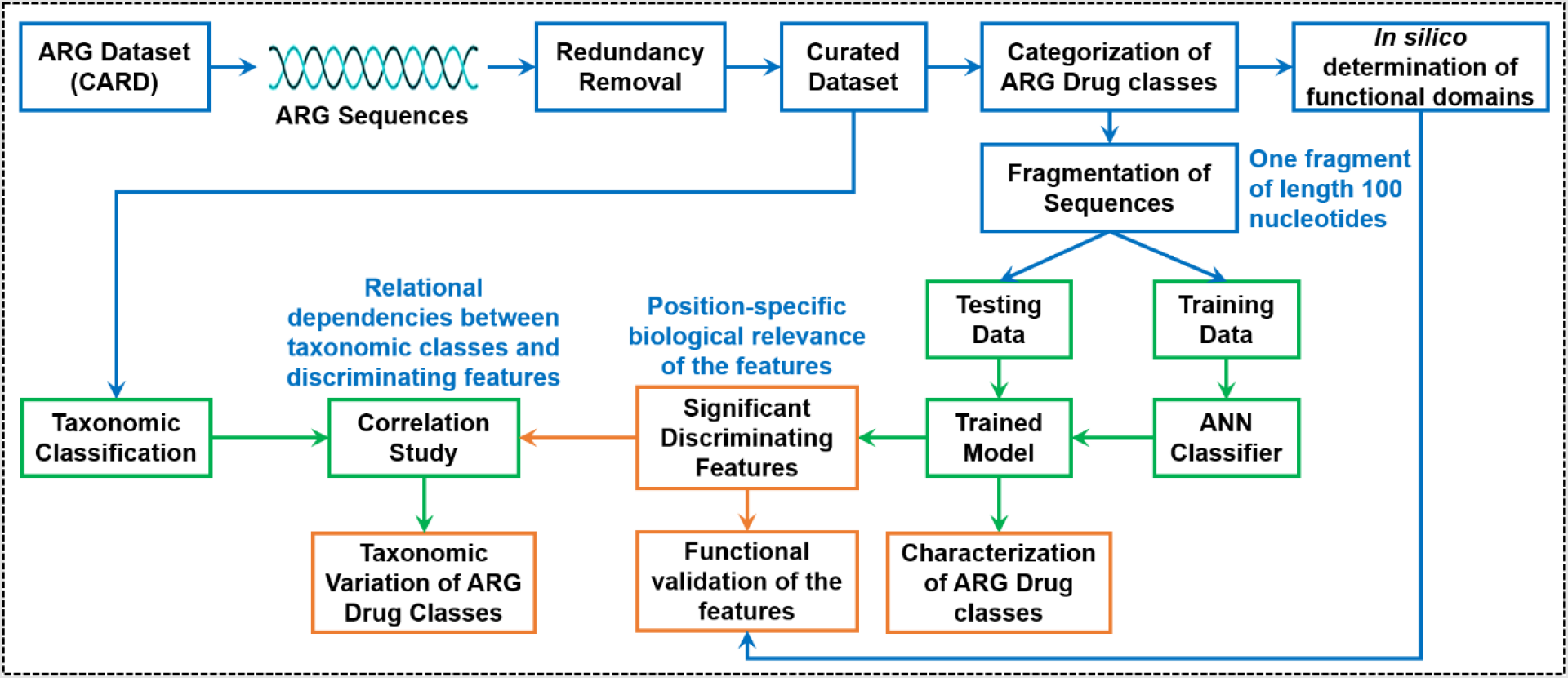
Overall process schematic for MACI

### 2.1. Dataset Curation and Pre-processing

From CARD (October 2021 release), we retrieved 2979 ARGs which were classified into 116 different drug classes. One drug class shows resistance towards a single antibiotic or multiple antibiotics. Drug classifications are vital as they defend us from severe side effects and confirm that our body can use the medicine properly [36, 37]. The drug classes are one of the imperative bases of Antibiotic Resistance Ontology (ARO) – the prime ontology in CARD. A drug class is a set of antibiotics with certain resemblances. Three central criteria are used to classify them viz., analogous chemical structures, similar mechanism of action and associated mode of action, and physiologic consequence i.e., how our body acknowledge them. CARD has organized ARO into three key branches – Determinant of Antibiotic Resistance, Antibiotic Molecule and Mechanism of Antibiotic Resistance. Each of these 2979 ARGs are provided with unique ARO numbers and they belong to a certain drug class [38].

Initial dataset with 2979 ARG sequences were first processed by CD-HIT-EST (http://weizhong-lab.ucsd.edu/cd-hit/) with the parameters “-c 0.9 -n 5” to cluster identical sequences. 1051 clusters were resulted, each representing a redundant set of genes from different drug classes. Within these 1051 clusters, 875 clusters represented only one single ARG sequence. Therefore, all of these 875 sequences were included in our data, as they were unique. The rest of the 176 clusters contained multiple ARG sequences. From these 176 clusters, we took one representative sequence from the 100% identical sequences, belonged to same drug class. We found such 2085 sequences. Consequently, completely identical genes from same or different drug-class were removed. The resulting set of 2960 ARG sequences could reflect inter-as well as intra drug-class diversity [39, 40].

Prior to the training, the curated database was analysed to check the distribution of class-specific sequence abundance. Each sequence was first divided into multiple fragments of equal length using a sliding window mechanism, having a stride of 1 nucleotide. The fragment length was optimized with respect to the performance of MACI and is discussed in section 3.1. Besides, while analysing the distribution of such nucleotide-fragments for 116 drug classes, we found that several classes comprise very low number of fragments (<8K) compared to highly sequence-populated drug classes (fragments >30K). Fig. 2(A) shows the abundance of the nucleotide fragments (in descending order) among various drug classes present in the CARD. We indexed the drug classes as C1, C2 … C116 maintaining the descending order. Detailed name of these representative drug classes is presented in Table 1 of supplementary material. Fig. 2(a) clearly reflects that only 22 classes are having more than 30K fragments, whereas remaining 94 drug classes comprise of fragments <30K. Importantly, class C1 (carbapenem;cephalosporin;penam) consists of more than 0.5 million fragments which is significantly higher than other drug classes. This can create a data imbalance problem as the classes having higher number of nucleotide fragments can outpower the classes with lower number of sequence fragments during training of the network. To overcome such data imbalance, we divided the entire dataset into two subsets – subset 1 was formed (22 classes) with more than 30K fragments, and remaining 94 drug classes (<30K fragments) created subset 2. Percentage contributions to the entire dataset for these 22 (subset 1) and 94 (subset 2) drug classes are showed in Fig. 2(b) and 2(c).

**Fig. 2.**
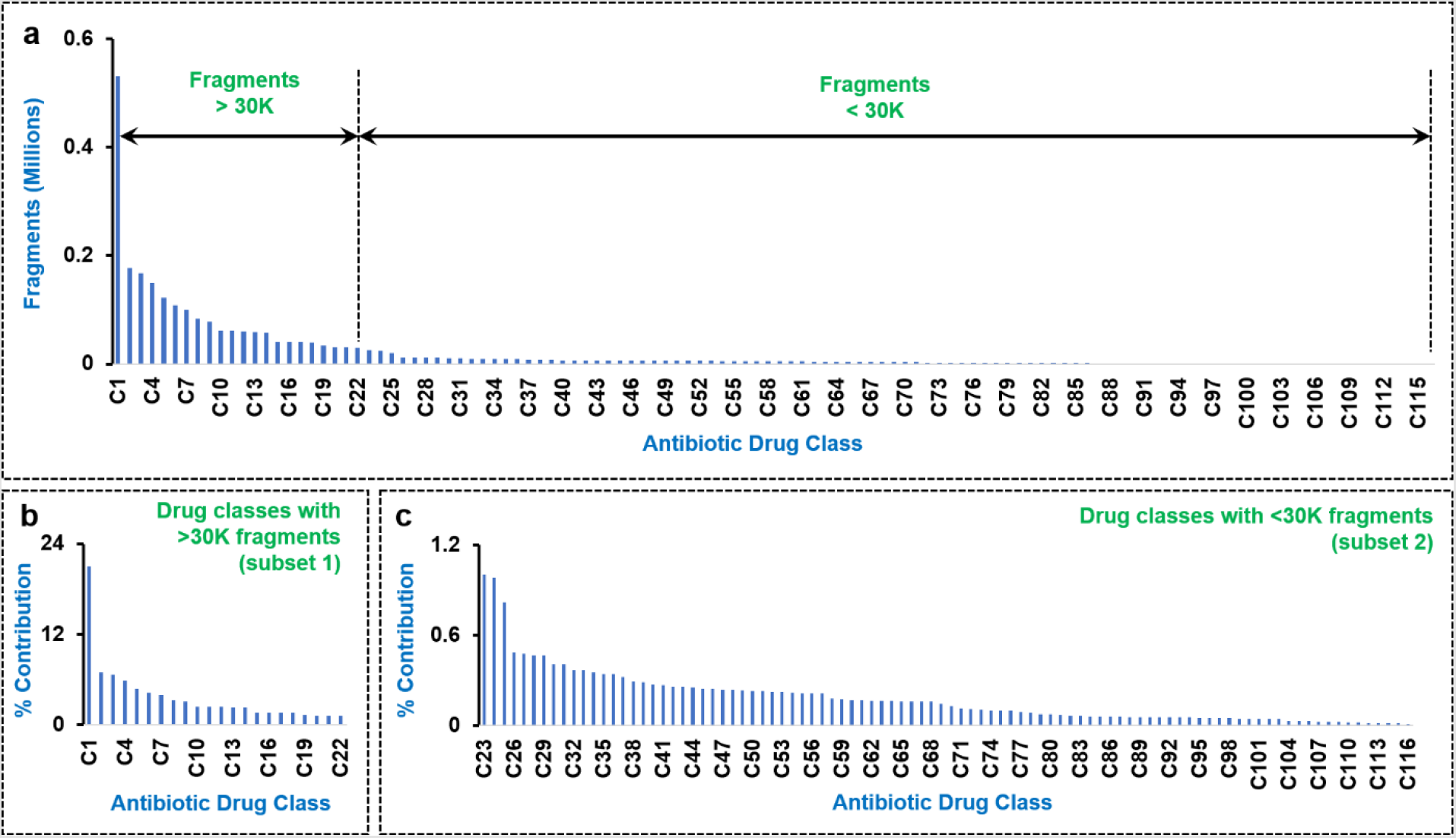
(a) Spread of number of fragments among 116 antibiotic drug classes. (b) Percentage contribution in overall data fragments of each class in subset 1. (c) Percentage contribution in overall data fragments of each class in subset 2.

As evident, class C1 holds a very strong number of fragments and contribute to 20.39% of the entire data. Apparently, contribution of other drug classes from subset 1 is large enough, whereas drug categories from subset 2 reflect a very less percentage of contribution (maximum 1% to overall data). The machine learning framework was trained on these two subsets (1 and 2) independently to reduce the tendency of underfitting. Final prediction of drug class for any unknown sequence was programmed to be the best output value among these two models.

### 2.2. Computational Framework

It should be noted that a brute search-based method to find the drug class of ARGs from a nucleotide fragment may lead to a huge computational time. This can be drastically reduced if a machine learning-based model is implemented. Here, we intend to develop an ANN framework for characterization of drug classes based on a sequence fragment. Prior to the implementation of machine learning model, the nucleotide fragments were encoded into a string of numerical values. We experimented with two encoding techniques – one-hot encoding and normalized label encoding. One-hot encoding automatically represented the input sequence into a matrix with binary values which was then fed to the input of the neural network. Other hand, normalized label encoding assigned a unique numerical value to each of the four nucleotides. The categorical data, containing a stream of A, C, T, G were manually encoded to a normalized numerical value. We provided a random order as C=1, T=2, G=3, A=4 and then normalized them as C=¼=0.25, T=½=0.5, G=¾=0.75, A=4/4=1. A comparative evaluation between these two was performed in terms of classification performance and is discussed in section 3.1.

A comprehensive block diagram of the computational framework is illustrated in Fig. 3, whereas the detailed neural network architecture is depicted in Fig. 1 of supplementary file. Post to the creation of the two subsets with 22 (subset 1) and 94 (subset 2) drug classes respectively, the same ANN-based model was trained independently using the data from each of the subsets. Therefore, two autonomous subset-specific ANN frameworks were modelled and optimized. This facilitated proper learning of the hidden patterns for the two subsets and overcome the underfitting issues due to data imbalance problem. Datasets for both the subsets were divided into training (80%) and testing (20%) groups with random sampling. For every subset, each sequence fragment was fed to the input of an ANN-based supervised learner, governed by backpropagation algorithm [41]. The number of input layer neurons were different for the two encoding techniques – to be accustomed with string data (in case of normalized label encoding) and binary matrix data (in case of one-hot encoding). Accordingly, the number of hidden layer neurons were varied as well. Following to the input layer of neurons, a densely packed network in hidden layers were designed. The number of neurons in these hidden layers were optimized against classification performance of the model (discussed in section 3.1). Number of neurons in the output layer was kept at 22 (for subset 1 drug category) and 94 (for subset 2 drug category). In this network architecture, the outputs of the previous layer were first linear summed and then passed through an activation function. Instead of using sigmoid or hyperbolic tangent rule as activation function in hidden layers, we implemented a rectified linear unit (ReLU). This can confront both linear and non-linear behaviour of the data, based on circumstances [42]. In general, both sigmoid and hyperbolic tangent activation functions are non-linear in nature and are preferred as they can fit more complex structures. Since, the proposed ANN-based framework was developed following the principle of backpropagation, the amount of error decreases as it passes through each layer and a derivative of the activation function was used to update the values. Therefore, use of either sigmoid or hyperbolic tangent activation functions lead to vanishing gradient problem where the updated values become so miniscule that they no longer impact the learning. Unlike to this, for ReLU, negative values were set to 0 and the rest were kept as the input itself. In terms of positive values, they were scaled and for negative values they were also set in a linear manner [42].

**Fig. 3.**
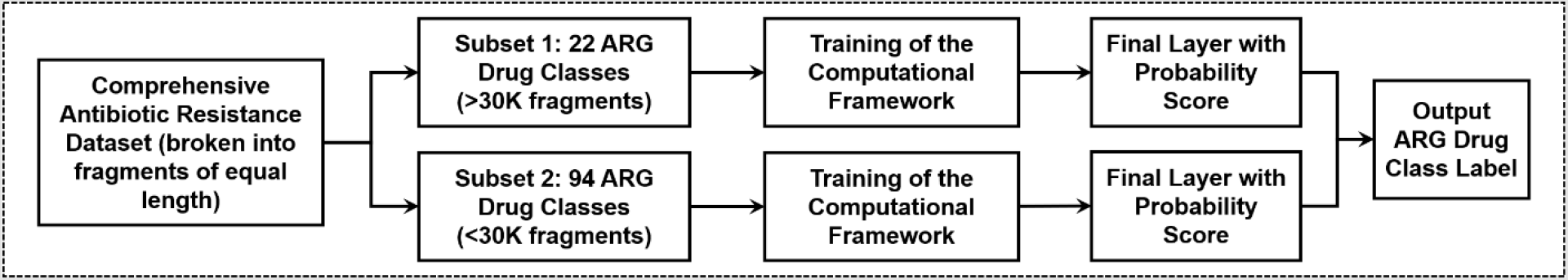
Framework for computational training of the MACI

Post to this, the network weights and bias were regularized using an optimizer. Since, the learning rate of optimizer is eventually a hyperparameter, it enables accurate training of the weights and bias. Among various notable optimization algorithms viz. stochastic gradient descent (SGD), root mean squared propagation (RMSPROP), adaptive gradient descent (AdaGrad), adaptive moment estimation (ADAM) etc., we adopted ADAM technique to augment the proposed computational model. In SGD, instead of considering the full data, the algorithm creates a batch of random samples and finds the minimal for each sample. This process is repeated through multiple epochs until it converges. Since, it explores through samples, the number of epochs required to find the minima is higher. Other hand, AdaGrad runs using an adaptive rule for changing the learning rate in each iteration. However, such changes are monotonic and aggressive which often lower the learning rate to a level that can compromise the accuracy of the model. Besides, RMSPROP makes use of the signs of the gradients, penalizing the main feature for incorrect classification. To overcome these limitations, ADAM explores exponentially weighted averages of the gradients to achieve faster training [43]. Unlike the RMSPROP technique which uses first moment, ADAM optimizer conceptualizes second moment, reducing the computational time to optimize network weights. The weighted averages were computed using,

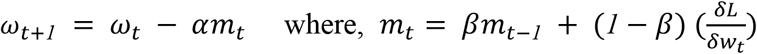

Here, *m*_*t*_ represents gradients aggregate at time t (initially, *m*_*t*_ = 0), *m*_*t* −*1*_ is gradients aggregate at time (t – 1), *ω*_*t*_ is the weights at time t, *ω*_*t* +*1*_ is weights at time (t + 1), α stands for learning rate at time t, δ*L* is derivatives of loss function, *δω*_*t*_ represents derivatives of weights at time t, and β is the moving average parameter. A SoftMax layer was added at the final output layer to represent the class labels as probability scores instead of a flat numeric [44]. This enabled a more generalized way of representing such multiclass classification. The class that matched for the given input sequence fragment, showed highest probability score. This represented an accurate measurement indicator of the model on the classification task.

Once properly trained, remaining 20% of the fragment-data from each of the subsets were utilized for testing of the computational framework. During characterization of the drug class of an unknown ARG sequence fragment, it was initially encoded and given as an input to both ANN models (for subset 1 and subset 2) independently. Based on the network weights, each of these models predicted a probability score corresponding to every drug class. Highest scoring value represented the drug class of the unknown ARG sequence.

Thereafter, we acquired class-specific common overlapping patterns which led to classification of an input fragment data. The model showcased a set of significant patterns of nucleotides for each sequence. These were further analysed using a separate in-house Python script to isolate class-specific common overlapping patterns. However, a question remains to be answered – whether the learning of the model had any biological significance or not. For this reason, the positions of these overlapping patterns were retrieved using the sliding window information that was used earlier to train the model. Then, the biological significances of these patterns were identified using an in-house Perl script (discussed in the next sub-section).

### 2.3. Prediction and analysis of functional domains

We predicted the biological functional domains or motifs for all sequences corresponding to subset 1 (22 antibiotic drug classes) using the Perl-script ‘ps_scan.pl’, available in the PROSITE database (https://prosite.expasy.org/) [45, 46], and then using an inhouse Perl-script we extracted the positions of the predicted functional domains in all sequences [47]. Subsequently, we matched these positions with locations of the significant patterns, predicted by our model for different drug classes of ARGs.

## 3. Results and Discussion

### 3.1. Parameter optimization

#### Fragment length

We found that the performance of MACI is highly sensitive to fragment length. Fig. 2 of the supplementary material showcases the variations in performance (in terms of average precision, recall and F1 score) of MACI with respect to fragment lengths, ranging from 50 to 140bp. The model showed a very poor performance when we kept the fragment lengths between 50-80bp. With smaller number of nucleotides, the neural network was not able to properly train its network parameters. This caused increment in the number of false negative and false positive cases. We found that the model’s performance was increased sharply in proportion to the length size till the value reached to 100bp. For 100bp<length<130bp, the changes in the performance score were not so significant, indicating a good possible region to choose the length size. Above length>130bp, the performance started falling again. Since, MACI is primarily developed to target metagenomic reads, in which case the read-length often is less. Considering average Illumina read-length of 150bp, we chose a length of 100bp to accumulate sufficient number of fragments from each read. Eventually, this led to attain a good performance score.

#### Encoding Technique

In a comparative performance evaluation of the encoding techniques (one-hot and normalized label encoding), the proposed ANN-based model showed an overall average precision of 0.833 and a recall of 0.813 while using normalized label encoding, whereas the scores became 0.847 and 0.821 respectively when implemented with one-hot encoding technique for subset 1 data. The changes in performance scores were not significant when associating either of the encoding methods. However, one-hot encoding was found to be more computationally expensive. As the process coded the sequence fragments into individual binary matrices, internal computations and architecture became more complex. This led to an overall increment of the computational burden. Hence, we opted to associate normalized label encoding in our proposed model for downstream analysis.

#### Number of neurons in hidden layers

Considering a fragment length of 100bp, the number of neurons in consecutive three hidden layers were optimized against classification performance of MACI through different trials. Both normalized label encoding and on-hot encoding were opted to perform this test. Results of these trials are tabulated in Table 2 (for normalized label encoding) and Table 3 (for one-hot encoding) of the supplementary material. We found that with a combination of 100, 400 and 200 number of neurons in successive three hidden layers (for normalized label encoding), the framework showed better performance in terms of overall average accuracy. Similarly, a combination of 400, 1500 and 500 number of neurons in successive three hidden layers for one-hot encoding resulted higher performance score.

### 3.2. Performance of the classifier – Overall

Performance of the proposed ANN-based computational model was verified against the testing data from subset 1 and 2 using an k-fold cross validation approach. Classification results were compared with respect to their annotated drug class labels, as per CARD. Class-specific performances of the model for some of these drug types are given in Fig. 4(a). For the drug class C1: “carbapenem;cephalosporin;penam”, approximately 86% of the predicted data (precision) and 89% of the actual data (recall) were rightly classified with an f1-score of 0.88. Similarly, precision and recall were measured for the other classes like - C2: “cephalosporin” 0.86 and 0.88; C3: “cephamycin” 0.84 and 0.85; C4: “cephalosporin;monobactam;penam;penem” 0.98 and 0.87; C9: “fluoroquinolone antibiotic” 0.93 and 0.91; C22: “penam;penem” 0.98 and 0.86 etc. Class-specific scores for subset 1 drug categories is tabulated in Table 4 of the supplementary material. These results validated the efficacy of MACI toward accurately categorizing the drug classes belongs to subset 1. This is because of the high number of fragments in this subset which enabled accurate training of the network weights, facilitating proper identification of hidden patterns. Other hand, for subset 2, the overall average precision dropped down to 0.57 with a recall of 0.53 for first 9 drug classes, and even lower to other drug categories. Lack of data in these drug classes (subset 2) restricted accurate tuning of the network parameters, leading to the increment of misclassification error. Moreover, the computational model could not accurately identify the discriminating patterns from this group. However, as evident, the proposed model showed a significant accuracy towards characterizing subset 1 drug classes, having sufficient number of fragments.

**Fig. 4.**
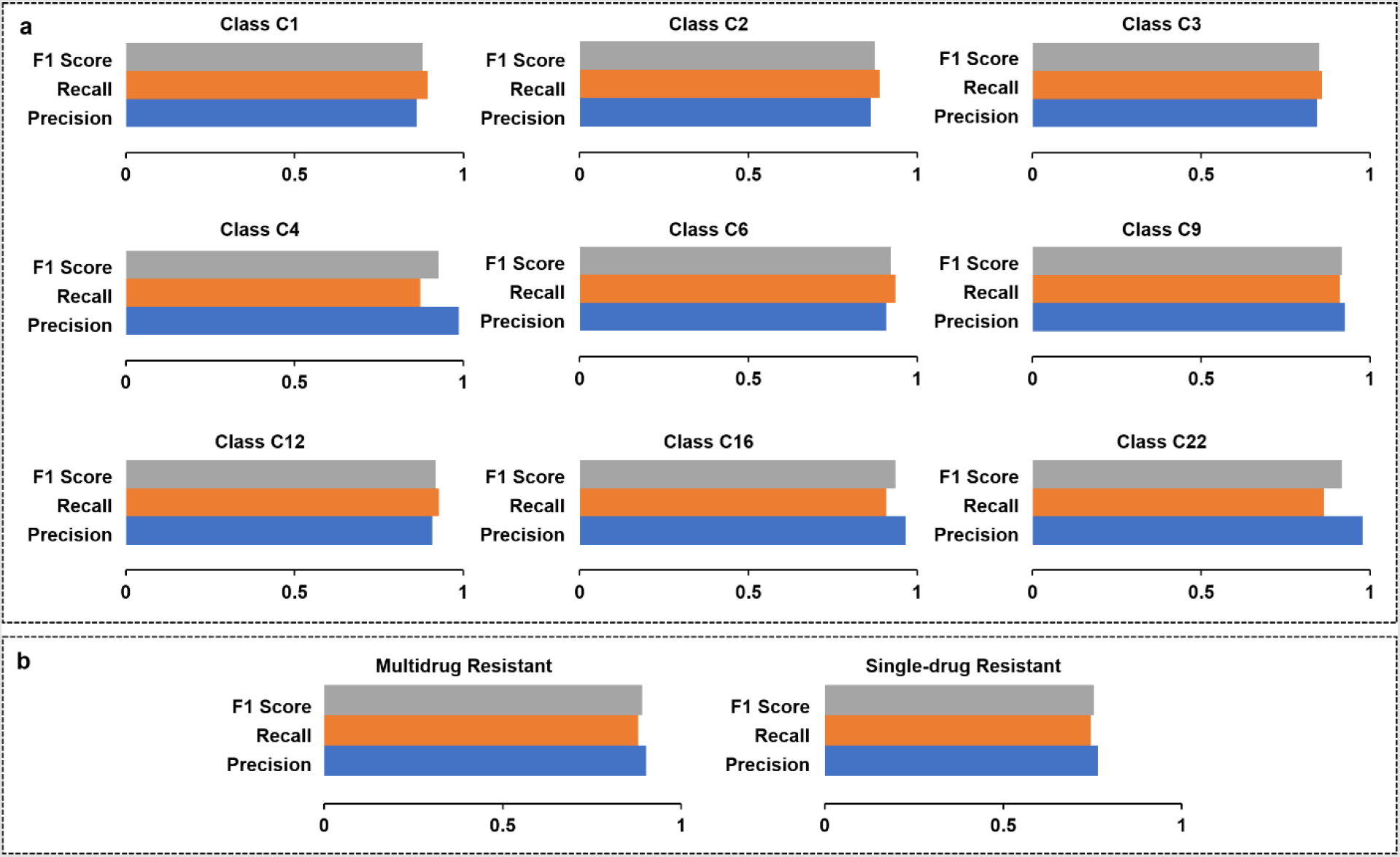
(a) Class-specific performance metrics of the proposed classifier model. (b) Performance evaluation of the proposed model – Multi-drug vs Single-drug resistant

### 3.3. Sensitivity analysis of k

In general, small value of k in k-fold cross validation technique leads to noisy estimate, whereas comparatively high value of k reduces the chances of noisy measurements. As for verification, we have performed a sensitivity analysis of k for three different values: k = 5, 8, and 10. For subset 1 data, average accuracies of the model were achieved to be 0.7852, 0.8264, and 0.8271 respectively for these three settings of k. As evident, both k = 8 and k = 10 can be used to obtain good performance of the model.

### 3.4. Performance of the classifier – multidrug vs single drug resistance

While analysing performance of the proposed model in the context of multidrug vs single-drug resistance for subset 1 data, we found that multidrug resistant categories were classified with higher precision and recall (0.9 and 0.88) in comparison to the single-drug resistant classes (0.77 and 0.75), as shown in Fig. 4(b). This reason is - the computational model characterized the drug classes based on a set of patterns. In these, we found multiple number of such patterns belonged to multidrug resistant categories, while one or two were contributed from single-drug resistant classes. This led to the subsequent reduction in false positive and false negative numbers for the multidrug resistant classes, triggering the increment in performance score.

### 3.5. Identification of class-specific patterns

Within 22 drug classes from subset 1, our model identified a set of nucleotide patterns from each sequence. These were further analysed to isolate class-specific overlapping patterns (using a separate in house Python script) respective to five drug classes viz. C1 (carbapenem;cephalosporin;penam), C2 (cephalosporin), C3 (cephamycin), C4 (cephalosporin;monobactam;penam;penem) and C9 (fluoroquinolone antibiotic). It should be noted that C1 and C4 are multidrug resistant whereas C2, C3, and C9 are single-drug resistant. To study the biological meaningfulness of these class-specific patterns, we mapped their positions within the biological functional domains for all sequences present in these classes (summary is provided in Table 5 of the supplementary material).

Analysis of these class-specific nucleotide signatures showed that there were 11864 sequence fragments from 723 ARG sequences in C1. These signatures were AAGC, GCAA, AAGCAA, GCAAGC and TGGA. AAGC and GCAA were observed to be more frequent than others. Three sequence patterns viz., AAGC, GCAA and TGGA (AAGC was most frequent) were found within two functional domains of 240 ARGs. One of the functional domains was “β-lactamase class-A active site” (PROSITE domain id: PS00146) which contained AAGC and GCAA. The other one was “β-lactamases class-B signature-2” (PROSITE domain id: PS00744), containing TGGA (as shown in Fig. 5). In general, β-lactamase enzymes catalyse hydrolysis of an amide bond in the β-lactam ring of cephalosporin antibiotics [48]. Among the four types of β-lactamase, class-A enzymes are serine hydrolases [49, 50]. Other hand, β-lactamases class-B signature-2 are able to hydrolyse carbapenem compounds [51, 52]. The action mechanism of C1 is “antibiotic inactivation” i.e., enzymatic inactivation of antibiotic to confer drug resistance [53]. Therefore, the functional domains, which contained C1-specific nucleotide patterns, sustained the action mechanism of C1. This resembled a strong class-specific biological importance of these nucleotide signatures.

**Fig. 5.**
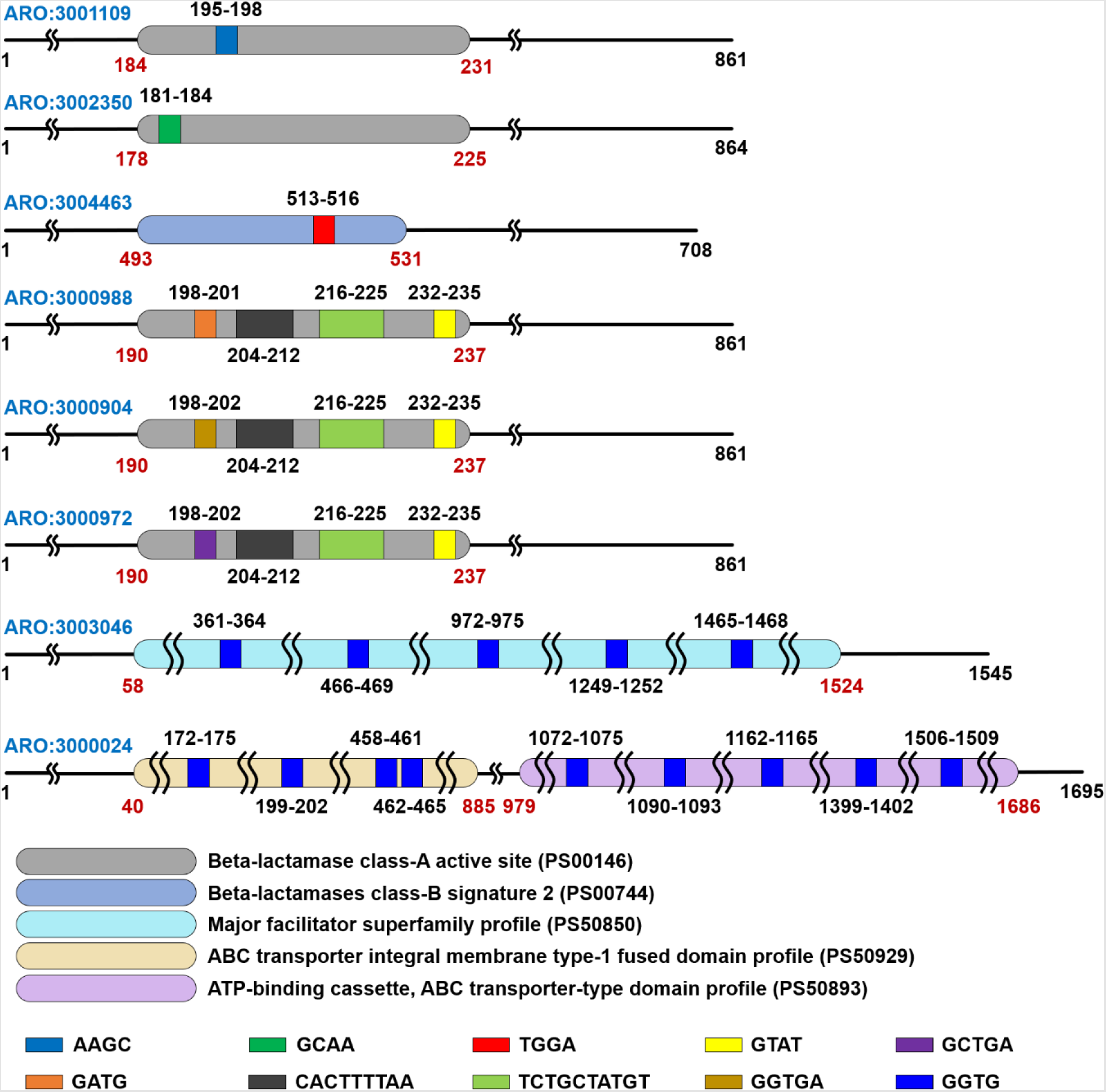
Pictorial depiction of the distribution of functional domains and sequence features for drug class C1 (ARO # 3001109, 3002350, 3004463), C4 (ARO # 3000988, 3000904, 3000972), and C9 (ARO # 3003046, 3000024)

For C4, there were 24957 sequence fragments from 196 ARG sequences. Although, these sequences were highly variable, still we got 62 C4-specific overlapping patterns. Among these, six patterns resided within the biological functional domains of nearly all the antibiotic resistance genes (195 out of 196). These nucleotide signatures were GATG, GTAT, GGTGA, GCTGA, CACTTTTAA and TCTGCTATGT (most frequent signatures were found to be CACTTTTAA, TCTGCTATGT, GATG and GTAT). All these six patterns were observed to be present within the functional domain “β-lactamase class-A active site”, as shown in Fig. 5. As discussed earlier, the action mechanism of C4 is also “antibiotic inactivation”. Therefore, the location of the C4- specific sequence patterns supported their biological relevance.

While analysing the single drug resistant class C2, we found 945 fragments for 206 ARG sequences. Although, the model predicted only one C2-specific pattern TGGT, we did not find its presence within the functional domains. In case of C3, we obtained 1262 fragments for 161 ARG sequences. Our model predicted only one signature TACT which was also not found within the functional domains. Similarly, for C9, there were 675 fragments from 119 ARG sequences. Our model predicted a nucleotide pattern GGTG which was found within functional domains of 14 ARGs in this drug class. GGTG was found to be present in several regions in three different functional domains viz., “Major facilitator superfamily (MFS)” (PROSITE domain id: PS50850), “ABC transporter integral membrane type-1 fused domain” (PROSITE domain id: PS50929) and “ATP-binding cassette, ABC transporter-type domain” (PROSITE domain id: PS50893) [54]. In general, there are two action mechanisms of C9. First is “reduced permeability to antibiotic” i.e., reduction in permeability to antibiotic due to reduced production of porins that is responsible for resistance [55, 56]. This is attained by the functional domain “Major facilitator superfamily (MFS) profile”. Since, MFS is one of the largest transporter families, it resides ubiquitously in almost all organisms. Second action mechanism is “antibiotic efflux” i.e., antibiotic resistance via out of cell transport of antibiotics [53]. Both “ABC transporter integral membrane type-1 fused domain” and “ATP-binding cassette, ABC transporter-type domain” help in this action. ABC transporters belong to the ATP-Binding Cassette (ABC) superfamily which utilizes hydrolysis of ATP for energising the varied biological import and export systems. Therefore, the positions of C9-specific pattern GGTG were found to be biologically relatable.

### 3.6. Analysis of taxonomic distribution for different drug classes

Additionally, we identified the taxonomic clades present in the above mentioned 5 antibiotic drug classes. As shown in Fig. 6, drug classes viz., C1, C2, C3, C4 and C9 showed the presence of 30, 18, 12, 19 and 18 bacterial genera respectively within them. *Acinetobacter, Aeromonas, Bacillus, Citrobacter, Enterobacter, Escherichia, Klebsiella, Proteus, Pseudomonas*, and *Salmonella* were found to be more frequent taxonomic classes. Three out of the above mentioned six core taxonomic clades viz., *Acinetobacter, Enterobacter* and *Klebsiella* (found in all the five drug classes) belong to ESKAPE pathogens. Their multidrug resistance property is quite well-known [47]. Other hand, the multidrug resistant strains of the rest three core taxonomic classes viz., *Citrobacter, Escherichia* and *Salmonella* are also causing clinical trouble worldwide. Therefore, categorising sequence-specific patterns of the ARGs present in these core taxonomic clades is of utmost importance for their easy and fast identification from metagenomic pools.

**Fig. 6.**
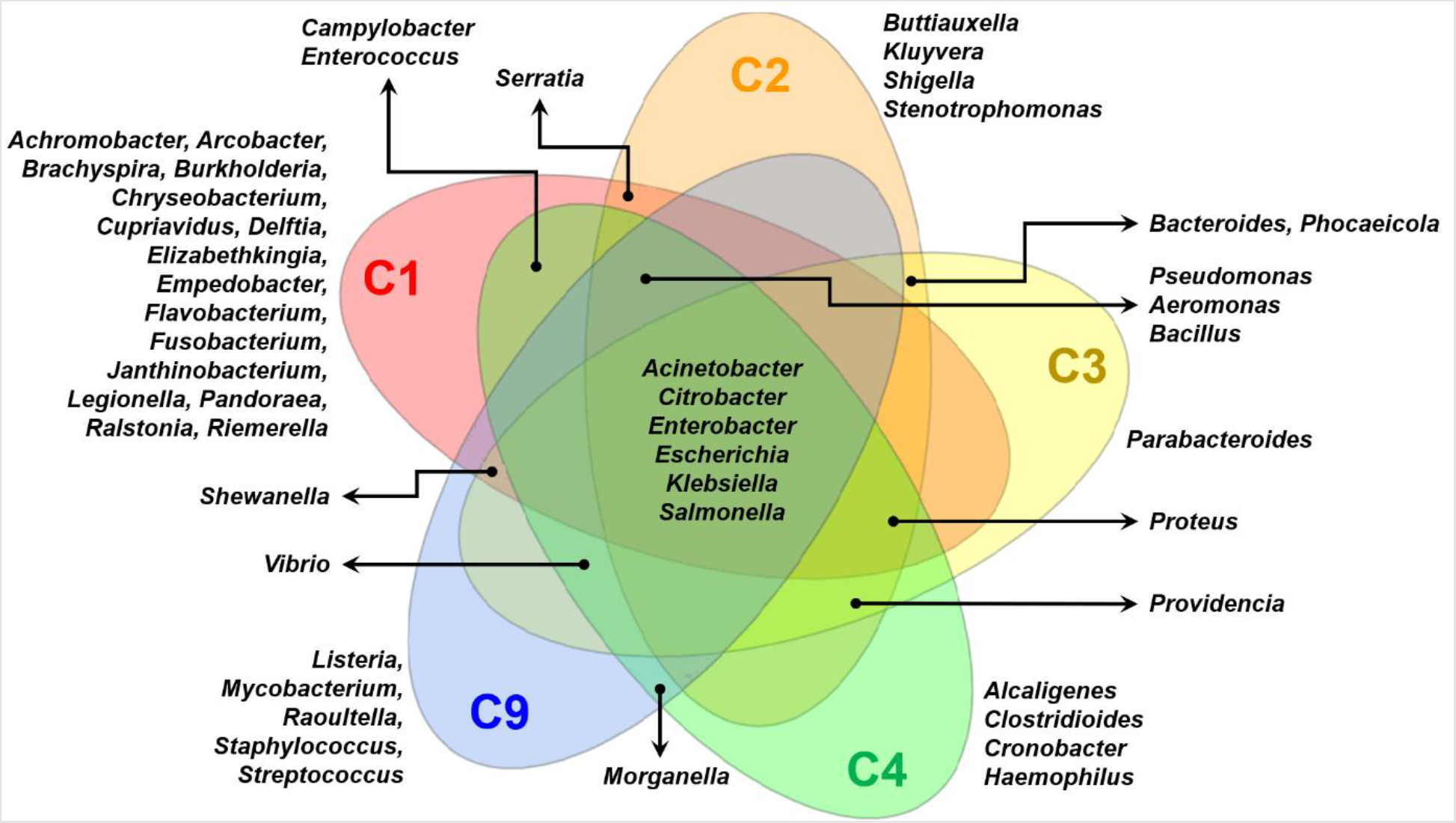
Taxonomic variations in antibiotic drug classes from subset 1

*Vibrio* and *Providencia* were observed to be the next frequent species. *Vibrio* was found in C3, C4 and C9 whereas *Providencia* was found in C2, C3 and C4. In case of *Vibrio* infection, usually drinking plenty of liquid to replace the fluid loss is necessary to treat the patients. However, in severe cases, antibiotics like quinolones, cephalosporins, tetracyclines, and penicillin are used. In this context, various studies explored *V. vulnificus* and *V. parahaemolyticus* as multiple antibiotic resistant. It was found that both environmental and clinical *Vibrio* isolates expressed similar kind of antibiotic resistance profiles [57, 58]. Other hand, fluoroquinolones, aminoglycosides, cephalosporins and β-lactam antibiotics are effective against *Providencia*. Being a life-threatening nosocomial pathogen, *Providencia* species (*P. stuartii* and *P. rettgeri*) were found to be resistant to multiple antibiotics [59].

Taxonomic clades, found in any two drug classes, were as follows – *Serratia* (C1 and C2), *Campylobacter* (C1 and C4), *Enterococcus* (C1 and C4), *Shewanella* (C1 and C9), *Bacteroides* (C2 and C3), and *Morganella* (C4 and C9). *Enterococcus* is a part of ESKAPE pathogens and is well-known for its multidrug resistance property. In general, *Campylobacter* infections are mild, but can be fatal for immuno-compromised individuals and children. *Bacterroides* (including *Phocaeicola*) infections are also associated with mortality. *Morganella* is a prominent one among the nosocomial infection causing bacterial species. *Serratia* and *Shewanella* are pathogenic in rare cases, but often show antibiotic resistance properties.

Furthermore, C1-specific taxonomic clades were *Achromobacter, Arcobacter, Brachyspira, Burkholderia, Chryseobacterium, Cupriavidus, Delftia, Elizabethkingia, Empedobacter, Flavobacterium, Fusobacterium, Janthinobacterium, Legionella, Pandoraea, Ralstonia* and *Riemerella*. C2-specific taxonomic clades were *Buttiauxella, Kluyvera, Shigella* and *Stenotrophomonas*. C3-specific taxonomic clade was only *Parabacteroides*. C4-specific taxonomic clades were *Alcaligenes, Clostridioides, Cronobacter* and *Haemophilus*. C9-specific taxonomic clades were *Listeria, Mycobacterium, Raoultella, Staphylococcus* and *Streptococcus*. C1 and C3 were found to be taxonomically most variable and conserved respectively. One probable reason for this is the widespread use of β-lactam antibiotics because of their selectivity, flexibility and low toxicity [60]. The cephalosporins, carbapenems, monobactams and penem are referred as β-lactam antibiotics. It should be noted that carbapenem resistance is a prevalent global public health threat [61]. Consequently, extensive use of cephalosporins can cause regular bacterial resistance to these drugs [62, 63]. The penam resistance is found in both gram-positive and gram-negative bacteria which also sustains higher taxonomic diversity for the penam-resistant organisms. Other hand, cephamycins are also β-lactam antibiotics with high similarity to cephalosporins. Less taxonomic diversity was found for cephamycin-resistant drug class (C3) due to recent emergence of cephamycin-resistant strains [64].

### 3.7. Validation of MACI using metagenomic data

To validate efficiency of the proposed model, we used faecal metagenome of 4 healthy samples from American individuals, available at National Center for Biotechnology Information (NCBI) Sequence Read Achieve (http://www.ncbi.nlm.nih.gov/sra) under the accession number ERP013933, BioProject number PRJEB12449 [65]. Metadata of these 4 samples includes health status, age, sex, and body-mass index. Sequencing data were generated using pair-end shotgun sequencing by Illumina technology [66]. After analysing these samples with MACI, we found the presence of aminoglycoside, tetracycline, carbapenem, cephalosporin, cephamycin, penam, fluoroquinolone, glycopeptide, lincosamide, macrolide, phenicol – with only a few data remained unclassified. Among these drug classes penicillin derivatives (penams), cephalosporins and cephamycins (cephems), monobactams and carbapenems belong to β-lactam antibiotics. In addition, macrolide and lincosinamide show similar mode of action i.e., they act as bacteriostatic agents by blocking bacterial protein synthesis, after binding to their 50S subunit. The abundance of each drug class was calculated using the sum of normalized values of the respective drug class. A comparative statistical analysis among these drug classes revealed the presence of five most abundant drug classes in our dataset, as shown in Fig. 7. They were aminoglycoside, beta-lactam, tetracycline, multidrug resistance and macrolide-lincosinamide. These five drug classes, having relatively higher abundance values, were further used for statistical analysis using student’s t-test. Eventually, these drug classes showed a p-value <0.05.

**Fig. 7.**
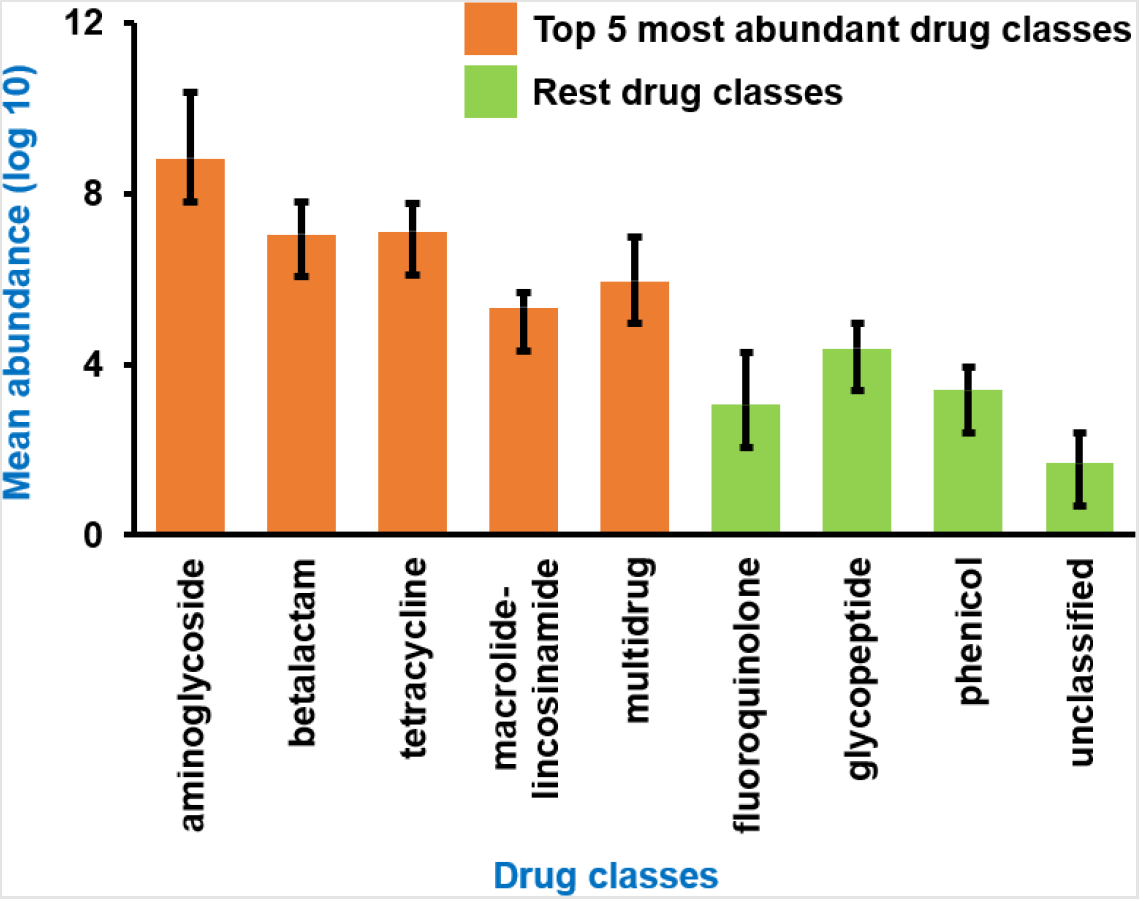
Abundance of different antibiotic drug classes detected by MACI

Earlier Qiu et al. used 2037 faecal metagenomes from 12 different countries and quantified the ARGs in each community [67]. The 4 healthy samples, used for the validation of MACI in this work, were also a part of the data utilized by Qiu et al. They found tetracycline, aminoglycoside, β-lactam, macrolide-lincosamide-streptogramin (MLS), and vancomycin resistance genes as dominant ARG types in the human gut. These ARG types were also observed by MACI, except the streptogramin. Drug classes containing streptogramin (C17 and C21 in subset 1) were multi drug resistant i.e., the ARGs in these drug classes showed resistance to several other antibiotics besides streptogramin. This might be a reason for not detecting specifically streptogramin during validation of MACI. In subset 2, C41 represented streptogramin as a single drug resistance and C52, C71, and C101 as multi drug resistance. These reflected a very less contribution of sequence fragments, leading to much lower accuracy in categorizing streptogramin antibiotics. Moreover, MACI identified three drug classes viz., fluoroquinolone, glycopeptide and phenicol in low rates. It should be noted, C11 drug class (in CARD) contains glycopeptide antibiotic. In general, first-generation glycopeptide antibiotics include vancomycin, teicoplanin, and ramoplanin. Detection of glycopeptide supports the presence of vancomycin as well. Other hand, fluoroquinolone and phenicol were not detected in the work done by Qiu et al. [67]. Additionally, Qiu et al. defined carbomycin, fusidic-acid, spectinomycin, puromycin, tetracenomycin C and fusaric-acid as rare ARGs – which were neither detected by MACI as well.

For the American antibiotic resistome, a relative bias toward aminoglycoside and less rate of MLS was found by Qiu et al. Similar trend was perceived by MACI – abundance of aminoglycoside was found to be significantly higher (p-value < 0.05) than the rest of the drug classes, except β-lactam and tetracycline. Overall, the findings by Qiu et al. corroborated the outcome of MACI. From implementational view-point, Qiu et al. used ARGs-OAP v2.0 pipeline to acquire ARG profiles, which included a two-step protocol [68]. First, UBLAST was implemented to identify probable ARG sequences. Second, BLASTX (a similarity-based search algorithm) was used to align them against database. The UBLAST algorithm performed local alignments for searching a database for a given E-value threshold [69]. Here, the proposed model, MACI, was developed using an ANN-based algorithm for classification of ARG-specific drug classes. Results of these two processes showed a little bit variation due to the differences in the protocol. Still, sufficient overlaps between the outcomes ascertain the actuality of the results obtained using MACI.

### 3.8. Comparison with existing pipelines

In order to establish our proposed model as a potential tool for identification of ARG-specific drug classes, we compared its performance with two existing pipelines – ARG-HMD and fARGene. ARG-HMD follows a hierarchical multi-task deep learning-based methodology to annotate and categorize drug classes of ARGs. Other hand, fARGene is a well-known machine learning-based pipeline, backboned by hidden markov model algorithm. Apart from these tools, there are a few non-alignment-based pipeline present which can annotate fasta sequences. Since our aim was to compare performance against the CARD database (composed of fasta sequences), we selected the aforementioned two tools, which have the desired capabilities.

While performing a comparative analysis, we found ARG-HMD could not classify 444 sequences and misclassified 117 sequences to be “non-ARG” for subset 1 data. ARG-HMD showed an average precision and recall of 0.1949 and 0.9731 respectively. Although the recall score is comparatively higher, MACI outperformed ARG-HMD with a significant difference in average precision score. Moreover, comparison of MACI with ARG-HMD yielded some basic differences. We found the outcome of ARG-HMD to be of lower resolution – MACI provides exact drug class as output, whereas ARG-HMD provides a much broader classification of sequences. Multiple CARD drug classes such as “carbapenem;cephalosporin;penam”, “cephalosporin”, “cephalosporin;monobactam;penam;penem”, “cephamycin” belong to the broader class β-lactam. ARG-HMD can classify the broader group (here, β-lactam), whereas MACI can distinguish between all the sub categories. Other hand, fARGene misclassified on an average 31.08% of input sequences to be “non-ARG” across multiple runs. A drawback of fARGene is its dependency on ARG-specific class models. To run fARGene successfully, it requires the user to have preliminary knowledge about the class of a sequence. Precision score could not be calculated for fARGene as each class-specific run was independent of each other and had overlaps. However, fARGene resulted an average recall of 0.67 which is significantly lower than MACI (0.89). Hence, our proposed model can not only provide a good classification score (for subset 1 data), but also can provide a deep classification of drug classes.

Although, the developed computational model can classify subset 1 drug classes with significant accuracy, its performance was observed to be decreased drastically for other drug categories (from subset 2). With comparatively low amount of sequence data, the model accuracy dropped – which is likely to be a generic case of such kind of machine learning-based model. If properly trained with good amount of data, we anticipate that the model performance can increase. As evidenced, the proposed technique can be utilized as a potential tool to characterize different drug classes of ARGs corresponding to subset 1 (having large amount of data to accurately train the neural network). As future work, indeed it would be great to incorporate more data and frame the computational pipeline into a software-based platform through which the users can easily and efficiently characterize drug classes of ARGs for any unknown sequence.

## 4. Conclusion

We developed an ANN-based computational model, MACI, which was applied on CARD data for accurate classification of drug classes of ARGs. The model showed a significant performance in classifying 22 (being populated with sufficient amount of sequence data) of 116 drug classes. We also observed that multidrug resistant classes were classified with higher precision and recall compared to single-drug resistant classes. This is because the neural network characterized the drug classes based on a set of nucleotide patterns, in which multiple number of such patterns belonged to multidrug resistant categories. Only one or two patterns were contributed from single drug resistant classes. In a separate analysis, the overlapping patterns for five drug classes were found to be more explicit as they were present in all class-specific sequences. Detailed analysis of these five drug classes indicated non-random nature of these significant patterns. For two multidrug resistant classes (“carbapenem;cephalosporin; penam” and “cephalosporin;monobactam;penam;penem”) and one single drug resistant class (fluoroquinolone antibiotic), some of these nucleotide signatures were observed within the biological functional domain. Eventually, they conserved their respective drug class-specific action mechanism. Additionally, taxonomic analysis revealed carbapenem;cephalosporin;penam-resistance (β-lactam antibiotics) to be taxonomically most variable. This supports the fact of extensive use of β-lactam antibiotics worldwide, leading to much higher bacterial resistance to these drugs. Although cephamycins are also β-lactam antibiotics with high similarity to cephalosporins, more recent emergence of cephamycin-resistant strains might be a reason of the less taxonomic diversity for the cephamycin-resistant drug class. Six core taxonomic clades (found in all the five drug classes) were found to be highly important from pathogenic point of view, posing global clinical threats with their multidrug resistance properties. Therefore, characterizing the ARG sequence-specific patterns and recognizing them on the basis of those from metagenomic pools, can provide an alternative solution and can help in the extension of the lifetime of current antibiotics.

## Acknowledgement

RB and KB would like to acknowledge the Indian Council of Medical Research (ICMR) for granting this project work through Extramural Adhoc funding (RFC No. ISRM/Adhoc/38/2020-21 dated 16.12.2020).

