## Supplementary Files for "MACI: A machine learning-based approach to identify drug classes of antibiotic resistance genes from metagenomic data"

### MACI: A machine learning-based approach to identify antibiotic drug classes from metagenomic data and their distribution in taxonomic clades

### Supplementary Material

Table 1. Mapping of class-names to their associated class-indexes

| Class Index | Name of Antibiotic Drug Class |
|---|---|
| C1 | carbapenem;cephalosporin;penam |
| C2 | cephalosporin |
| C3 | cephamycin |
| C4 | cephalosporin;monobactam;penam;penem |
| C5 | aminoglycoside antibiotic |
| C6 | carbapenem;cephalosporin;monobactam |
| C7 | peptide antibiotic |
| C8 | tetracycline antibiotic |
| C9 | fluoroquinolone antibiotic |
| C10 | cephalosporin;penam |
| C11 | glycopeptide antibiotic |
| C12 | carbapenem;cephalosporin;cephamycin;penam;penem |
| C13 | carbapenem;cephalosporin;cephamycin;penam |
| C14 | penam |
| C15 | phenicol antibiotic |
| C16 | carbapenem;cephalosporin;monobactam;penam |
| C17 | lincosamide antibiotic;macrolide antibiotic;oxazolidinone antibiotic;phenicol antibiotic;pleuromutilin antibiotic;streptogramin antibiotic;tetracycline antibiotic |
| C18 | carbapenem |
| C19 | macrolide antibiotic |
| C20 | cephalosporin;cephamycin |

| | |
|---|---|
| C21 | lincosamide antibiotic;macrolide antibiotic;streptogramin antibiotic |
| C22 | penam;penem |
| C23 | cephalosporin;monobactam;penam |
| C24 | cephalosporin;monobactam |
| C25 | diaminopyrimidine antibiotic |
| C26 | fosfomycin |
| C27 | carbapenem;cephalosporin;cephamycin;monobactam;penam |
| C28 | cephalosporin;cephamycin;penam |
| C29 | glycylcycline;tetracycline antibiotic |
| C30 | lincosamide antibiotic |
| C31 | rifamycin antibiotic |
| C32 | aminoglycoside antibiotic;fluoroquinolone antibiotic |
| C33 | mupirocin |
| C34 | aminocoumarin antibiotic |
| C35 | aminocoumarin antibiotic;macrolide antibiotic;monobactam;tetracycline antibiotic |
| C36 | cephalosporin;cephamycin;monobactam;penam;penem |
| C37 | carbapenem;cephalosporin |
| C38 | aminoglycoside antibiotic;cephalosporin;cephamycin;penam |
| C39 | cephalosporin;fluoroquinolone antibiotic;glycylcycline;penam;phenicol antibiotic;rifamycin antibiotic;tetracycline antibiotic;triclosan |
| C40 | peptide antibiotic;rifamycin antibiotic |
| C41 | streptogramin antibiotic |
| C42 | acridine dye;disinfecting agents and intercalating dyes;fluoroquinolone antibiotic |
| C43 | fluoroquinolone antibiotic;tetracycline antibiotic |
| C44 | triclosan |
| C45 | carbapenem;cephalosporin;diaminopyrimidine antibiotic;fluoroquinolone antibiotic;lincosamide antibiotic;macrolide antibiotic;penem;phenicol antibiotic;rifamycin antibiotic;tetracycline antibiotic |
| C46 | macrolide antibiotic;penam |

| | |
|---|---|
| C47 | carbapenem;cephalosporin;cephamycin;monobactam;penam;penem;phenicol antibiotic |
| C48 | fluoroquinolone antibiotic;macrolide antibiotic;penam |
| C49 | cephalosporin;fluoroquinolone antibiotic;fusidic acid;macrolide antibiotic |
| C50 | carbapenem;cephalosporin;monobactam;penam;penem |
| C51 | acridine dye;disinfecting agents and intercalating dyes;fluoroquinolone antibiotic;tetracycline antibiotic |
| C52 | lincosamide antibiotic;oxazolidinone antibiotic;phenicol antibiotic;pleuromutilin antibiotic;streptogramin antibiotic |
| C53 | diaminopyrimidine antibiotic;fluoroquinolone antibiotic;phenicol antibiotic |
| C54 | acridine dye;carbapenem;diaminopyrimidine antibiotic;disinfecting agents and intercalating dyes;macrolide antibiotic;phenicol antibiotic;tetracycline antibiotic |
| C55 | aminocoumarin antibiotic;aminoglycoside antibiotic;cephalosporin;diaminopyrimidine antibiotic;fluoroquinolone antibiotic;macrolide antibiotic;penam;phenicol antibiotic;tetracycline antibiotic |
| C56 | fluoroquinolone antibiotic;macrolide antibiotic;phenicol antibiotic;tetracycline antibiotic |
| C57 | aminoglycoside antibiotic;cephalosporin;fluoroquinolone antibiotic;macrolide antibiotic |
| C58 | macrolide antibiotic;tetracycline antibiotic;triclosan |
| C59 | carbapenem;cephalosporin;cephamycin;monobactam;penam;penem |
| C60 | cephalosporin;penam;peptide antibiotic |
| C61 | acridine dye;disinfecting agents and intercalating dyes;nucleoside antibiotic |
| C62 | aminocoumarin antibiotic;carbapenem;cephalosporin;cephamycin;diaminopyrimidine antibiotic;fluoroquinolone antibiotic;macrolide antibiotic;monobactam;penam;penem;peptide antibiotic;phenicol antibiotic;sulfonamide antibiotic;tetracycline antibiotic |
| C63 | diaminopyrimidine antibiotic;fluoroquinolone antibiotic;glycylcycline;nitrofuran antibiotic;tetracycline antibiotic |
| C64 | acridine dye;aminoglycoside antibiotic;carbapenem;cephalosporin;cephamycin;disinfecting agents and intercalating dyes;fluoroquinolone antibiotic;macrolide antibiotic;penam;phenicol antibiotic;tetracycline antibiotic |
| C65 | cephalosporin;cephamycin;fluoroquinolone antibiotic;penam |
| C66 | fluoroquinolone antibiotic;macrolide antibiotic;penam;tetracycline antibiotic |
| C67 | acridine dye;disinfecting agents and intercalating dyes;fluoroquinolone antibiotic;macrolide antibiotic;phenicol antibiotic;tetracycline antibiotic |
| C68 | lincosamide antibiotic;macrolide antibiotic |
| C69 | fusidic acid |

| | |
|---|---|
| C70 | aminocoumarin antibiotic;aminoglycoside antibiotic |
| C71 | lincosamide antibiotic;oxazolidinone antibiotic;phenicol antibiotic;streptogramin antibiotic |
| C72 | nucleoside antibiotic |
| C73 | fluoroquinolone antibiotic;macrolide antibiotic;rifamycin antibiotic |
| C74 | aminoglycoside antibiotic;carbapenem;cephalosporin;fluoroquinolone antibiotic;macrolide antibiotic;penam;penem;peptide antibiotic |
| C75 | cephalosporin;fluoroquinolone antibiotic;monobactam |
| C76 | antibacterial free fatty acids |
| C77 | aminocoumarin antibiotic;carbapenem;peptide antibiotic;rifamycin antibiotic |
| C78 | sulfonamide antibiotic;sulfone antibiotic |
| C79 | carbapenem;cephalosporin;cephamycin |
| C80 | pleuromutilin antibiotic |
| C81 | acridine dye;aminoglycoside antibiotic;carbapenem;cephalosporin;cephamycin;disinfecting agents and intercalating dyes;fluoroquinolone antibiotic;macrolide antibiotic;monobactam;penam;penem;phenicol antibiotic;tetracycline antibiotic |
| C82 | nitroimidazole antibiotic |
| C83 | elfamycin antibiotic |
| C84 | cephalosporin;penam;penem |
| C85 | aminoglycoside antibiotic;fluoroquinolone antibiotic;rifamycin antibiotic;tetracycline antibiotic |
| C86 | isoniazid;rifamycin antibiotic |
| C87 | penam;tetracycline antibiotic |
| C88 | carbapenem;penam |
| C89 | aminocoumarin antibiotic;aminoglycoside antibiotic;carbapenem;cephalosporin;cephamycin;fluoroquinolone antibiotic;glycylcycline;macrolide antibiotic;penam;penem;peptide antibiotic;phenicol antibiotic;rifamycin antibiotic;tetracycline antibiotic;triclosan |
| C90 | acridine dye;disinfecting agents and intercalating dyes;macrolide antibiotic |
| C91 | acridine dye;aminocoumarin antibiotic;aminoglycoside antibiotic;carbapenem;cephalosporin;cephamycin;diaminopyrimidine antibiotic;disinfecting agents and intercalating |

| | |
|---|---|
| | dyes;fluoroquinolone antibiotic;macrolide antibiotic;monobactam;penam;penem;peptide antibiotic;phenicol antibiotic;sulfonamide antibiotic;tetracycline antibiotic |
| C92 | aminoglycoside antibiotic;diaminopyrimidine antibiotic;macrolide antibiotic;oxazolidinone antibiotic;phenicol antibiotic |
| C93 | aminoglycoside antibiotic;benzalkonium chloride;fluoroquinolone antibiotic |
| C94 | acridine dye;disinfecting agents and intercalating dyes |
| C95 | aminoglycoside antibiotic;carbapenem;fluoroquinolone antibiotic;macrolide antibiotic;tetracycline antibiotic |
| C96 | acridine dye;disinfecting agents and intercalating dyes;fluoroquinolone antibiotic;triclosan |
| C97 | fluoroquinolone antibiotic;nitroimidazole antibiotic;tetracycline antibiotic |
| C98 | fluoroquinolone antibiotic;macrolide antibiotic |
| C99 | acridine dye;disinfecting agents and intercalating dyes;fluoroquinolone antibiotic;lincosamide antibiotic;nucleoside antibiotic;phenicol antibiotic |
| C100 | benzalkonium chloride;rhodamine;tetracycline antibiotic |
| C101 | lincosamide antibiotic;macrolide antibiotic;streptogramin antibiotic;tetracycline antibiotic |
| C102 | bicyclomycin |
| C103 | acridine dye;disinfecting agents and intercalating dyes;fluoroquinolone antibiotic;nucleoside antibiotic;phenicol antibiotic |
| C104 | sulfonamide antibiotic |
| C105 | carbapenem;monobactam;penam |
| C106 | disinfecting agents and intercalating dyes |
| C107 | aminocoumarin antibiotic;aminoglycoside antibiotic;carbapenem;cephalosporin;cephamycin;diaminopyrimidine antibiotic;fluoroquinolone antibiotic;macrolide antibiotic;monobactam;penam;penem;peptide antibiotic;phenicol antibiotic;sulfonamide antibiotic;tetracycline antibiotic |
| C108 | cephalosporin;cephamycin;fluoroquinolone antibiotic;glycylcycline;penam;phenicol antibiotic;rifamycin antibiotic;tetracycline antibiotic;triclosan |
| C109 | carbapenem;cephalosporin;cephamycin;fluoroquinolone antibiotic;glycylcycline;monobactam;penam;penem;phenicol antibiotic;rifamycin antibiotic;tetracycline antibiotic;triclosan |
| C110 | aminoglycoside antibiotic;cephalosporin;macrolide antibiotic;peptide antibiotic;rifamycin antibiotic;tetracycline antibiotic |
| C111 | aminoglycoside antibiotic;phenicol antibiotic;tetracycline antibiotic |

| | |
|---|---|
| C112 | acridine dye;cephalosporin;disinfecting agents and intercalating dyes;fluoroquinolone antibiotic;glycylcycline;penam;phenicol antibiotic;rifamycin antibiotic;tetracycline antibiotic;triclosan |
| C113 | acridine dye;cephalosporin;disinfecting agents and intercalating dyes;fluoroquinolone antibiotic;penam;peptide antibiotic;tetracycline antibiotic |
| C114 | peptide antibiotic;polyamine antibiotic |
| C115 | cephalosporin;cephamycin;fluoroquinolone antibiotic;macrolide antibiotic;penam;tetracycline antibiotic |
| C116 | aminocoumarin antibiotic;macrolide antibiotic |

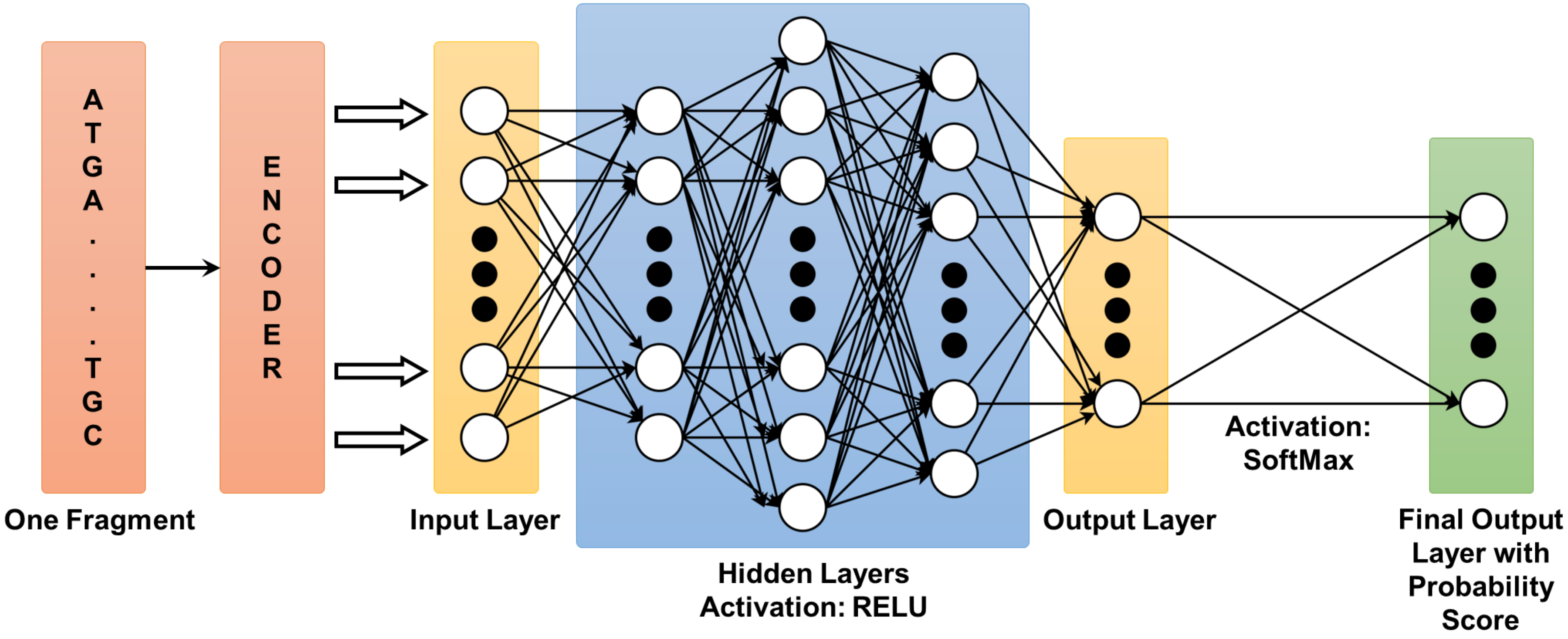


Fig.1. Architecture of ANN-based computational model. The number of neurons in each layer was different for the encoding techniques. For normalized label encoding – number of neurons in input and consecutive three hidden layers were finalized at 100, 100, 400, 200 respectively. For one-hot encoding – number of neurons in input and consecutive three hidden layers were finalized at 400, 400, 1500, 500. These numbers were optimized through various trials with respect to classification performance of the computational model, considering a fragment length of 100. Number of neurons in the output layer was kept at 22 (for subset 1 drug category) and 94 (for subset 2 drug category).

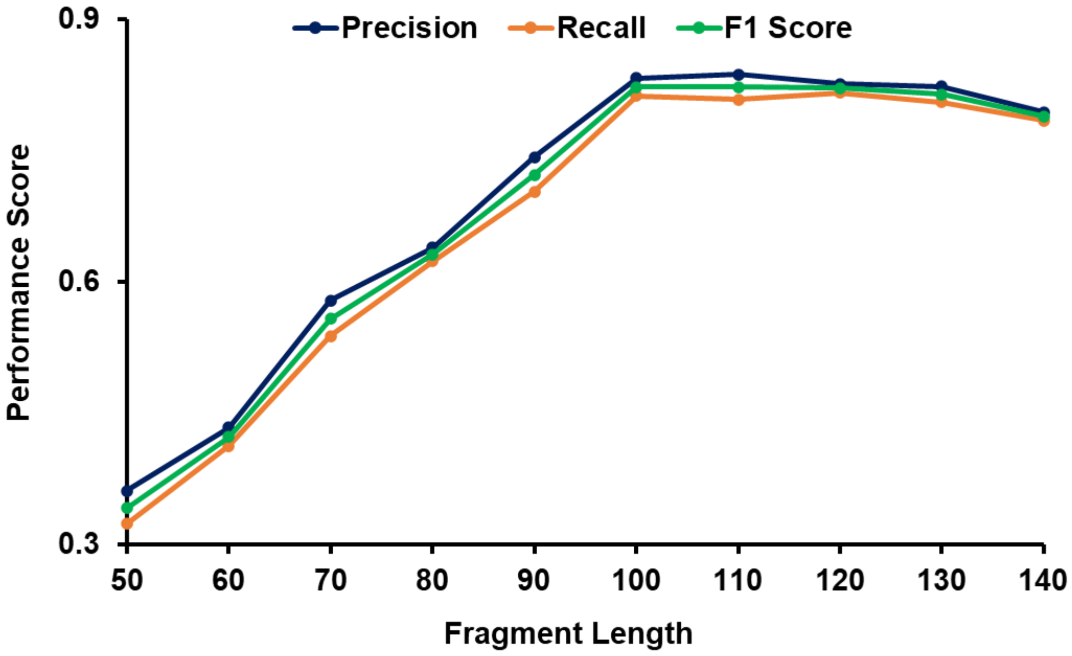


Fig. 2. Performance comparison of MACI (in terms of precision, recall and F1 score) with respect to different fragment lengths (using normalized label encoding). The model's performance was increased sharply in proportion to the length size till the value reached to 100bp. For 100bp<length<130bp, the changes in the performance score were not so significant, indicating a good possible region to choose the length size. Above length>130bp, we observed that the performance of MACI started falling again.

Table 2. Performance comparison (in terms of overall average accuracy) of the ANN framework with respect to number of neurons in successive three hidden layers – using normalized label encoding

| **Accuracy – with 50 neurons in hidden layer 1** | | | | | | |
|---|---|---|---|---|---|---|
| | | **Number of neurons in hidden layer 3** | | | | |
| | | **100** | **150** | **200** | **250** | **300** |
| **Number of neurons in hidden layer 2** | **300** | 0.35671 | 0.37238 | 0.43728 | 0.41268 | 0.36395 |
| | **350** | 0.38924 | 0.40924 | 0.47296 | 0.43172 | 0.39824 |
| | **400** | 0.44782 | 0.47293 | 0.57639 | 0.50374 | 0.47273 |
| | **450** | 0.41923 | 0.42395 | 0.46823 | 0.44703 | 0.41692 |
| | **500** | 0.35896 | 0.37479 | 0.42173 | 0.39274 | 0.36294 |

| **Accuracy – with 100 neurons in hidden layer 1** | | | | | | |
|---|---|---|---|---|---|---|
| | | **Number of neurons in hidden layer 3** | | | | |
| | | **100** | **150** | **200** | **250** | **300** |
| **Number of neurons in hidden layer 2** | **300** | 0.52951 | 0.55683 | 0.68914 | 0.64894 | 0.60945 |
| | **350** | 0.67925 | 0.68924 | 0.71763 | 0.69625 | 0.64397 |
| | **400** | 0.71364 | 0.74216 | **0.82638** | 0.77316 | 0.78762 |
| | **450** | 0.68581 | 0.69723 | 0.72536 | 0.71932 | 0.67291 |
| | **500** | 0.57392 | 0.62681 | 0.68741 | 0.67159 | 0.61943 |

| **Accuracy – with 150 neurons in hidden layer 1** | | | | | | |
|---|---|---|---|---|---|---|
| | | **Number of neurons in hidden layer 3** | | | | |
| | | **100** | **150** | **200** | **250** | **300** |
| **Number of neurons in hidden layer 2** | **300** | 0.42378 | 0.46927 | 0.52774 | 0.48753 | 0.44792 |
| | **350** | 0.49832 | 0.52262 | 0.58395 | 0.53466 | 0.50936 |
| | **400** | 0.58791 | 0.61837 | 0.68538 | 0.62914 | 0.58354 |
| | **450** | 0.52374 | 0.56495 | 0.61489 | 0.58426 | 0.54497 |
| | **500** | 0.47328 | 0.50194 | 0.54739 | 0.51387 | 0.49263 |

Table 3. Performance comparison (in terms of overall average accuracy) of the ANN framework with respect to number of neurons in successive three hidden layers – using one-hot encoding)

| **Accuracy – with 350 neurons in hidden layer 1** | | | | | | |
|---|---|---|---|---|---|---|
| | | **Number of neurons in hidden layer 3** | | | | |
| | | **400** | **450** | **500** | **550** | **600** |
| **Number of neurons in hidden layer 2** | **1400** | 0.35612 | 0.37834 | 0.44342 | 0.41817 | 0.37925 |
| | **1450** | 0.37372 | 0.42349 | 0.47773 | 0.42932 | 0.39278 |
| | **1500** | 0.45397 | 0.48581 | 0.58673 | 0.51853 | 0.48532 |
| | **1550** | 0.43072 | 0.41128 | 0.46326 | 0.45926 | 0.42371 |
| | **1600** | 0.36175 | 0.37924 | 0.41092 | 0.40735 | 0.36916 |

| **Accuracy – with 400 neurons in hidden layer 1** | | | | | | |
|---|---|---|---|---|---|---|
| | | **Number of neurons in hidden layer 3** | | | | |
| | | **400** | **450** | **500** | **550** | **600** |
| **Number of neurons in hidden layer 2** | **1400** | 0.53184 | 0.56382 | 0.69235 | 0.64936 | 0.59384 |
| | **1450** | 0.67134 | 0.69182 | 0.73284 | 0.68834 | 0.63829 |
| | **1500** | 0.72849 | 0.76823 | **0.83126** | 0.78267 | 0.77324 |
| | **1550** | 0.69392 | 0.71154 | 0.72925 | 0.70463 | 0.68382 |
| | **1600** | 0.59934 | 0.61873 | 0.67935 | 0.68673 | 0.61473 |

| **Accuracy – with 450 neurons in hidden layer 1** | | | | | | |
|---|---|---|---|---|---|---|
| | | **Number of neurons in hidden layer 3** | | | | |
| | | **400** | **450** | **500** | **550** | **600** |
| **Number of neurons in hidden layer 2** | **1400** | 0.43372 | 0.47733 | 0.53923 | 0.49382 | 0.45193 |
| | **1450** | 0.48447 | 0.53392 | 0.58218 | 0.53361 | 0.51472 |
| | **1500** | 0.60726 | 0.62832 | 0.70133 | 0.63924 | 0.60362 |
| | **1550** | 0.53924 | 0.55392 | 0.61738 | 0.59346 | 0.55394 |
| | **1600** | 0.48173 | 0.51628 | 0.53923 | 0.52448 | 0.49748 |

Table 4. Class-specific prediction scores of 22 ARG drug classes (belongs to subset 1)

| Class # | CLASS_NAME | PRECISION | RECALL | F1 Score |
|---|---|---|---|---|
| C1 | carbapenem;cephalosporin;penam | 0.862642741 | 0.8941 | 0.87808972 |
| C2 | cephalosporin | 0.862880886 | 0.8785 | 0.8706204 |
| C3 | cephamycin | 0.843580915 | 0.8475 | 0.84553592 |
| C4 | cephalosporin;monobactam;penam;penem | 0.984433013 | 0.8725 | 0.92597506 |
| C5 | aminoglycoside antibiotic | 0.738671875 | 0.700213 | 0.71892847 |
| C6 | carbapenem;cephalosporin;monobactam | 0.908737864 | 0.936 | 0.92216749 |
| C7 | peptide antibiotic | 0.723348801 | 0.7016375 | 0.71232775 |
| C8 | tetracycline antibiotic | 0.726600791 | 0.7028021 | 0.71450333 |
| C9 | fluoroquinolone antibiotic | 0.925653048 | 0.9061 | 0.91577216 |
| C10 | cephalosporin;penam | 0.845423729 | 0.8403 | 0.84285408 |
| C11 | glycopeptide antibiotic | 0.712230216 | 0.680134 | 0.69581217 |
| C12 | carbapenem;cephalosporin;cephamycin;penam;penem | 0.90739726 | 0.928 | 0.91758299 |
| C13 | carbapenem;cephalosporin;cephamycin;penam | 0.871292373 | 0.8225 | 0.84619342 |
| C14 | penam | 0.725384615 | 0.6907192 | 0.70762761 |
| C15 | phenicol antibiotic | 0.733687177 | 0.703765 | 0.71841466 |
| C16 | carbapenem;cephalosporin;monobactam;penam | 0.966471527 | 0.908 | 0.93632379 |
| C17 | lincosamide antibiotic;macrolide antibiotic;oxazolidinone antibiotic;phenicol antibiotic;pleuromutilin antibiotic;streptogramin antibiotic;tetracycline antibiotic | 0.920450014 | 0.9387 | 0.92948543 |
| C18 | carbapenem | 0.723300248 | 0.71585 | 0.71955584 |
| C19 | macrolide antibiotic | 0.709756098 | 0.7180025 | 0.71385548 |
| C20 | cephalosporin;cephamycin | 0.828954147 | 0.8045 | 0.81654402 |
| C21 | lincosamide antibiotic;macrolide antibiotic;streptogramin antibiotic | 0.831874203 | 0.8375 | 0.83467762 |
| C22 | penam;penem | 0.977916195 | 0.8635 | 0.91715348 |
| | **AVERAGE** | **0.833213079** | **0.813219241** | **0.822687217** |

Table 5. Summary of the drug class specific sequence features and the functional domains of the ARGs in which these features reside (Class indices are described in the supplementary table 1)

| | **Multi-drug resistant classes** | | | **Single-drug resistant classes** | | |
|---|---|---|---|---|---|---|
| **Class #** | C1 | | C4 | C2 | C3 | C9 |
| **Resistant to** | carbapenem;cephalosporin;penam | | cephalosporin; monobactam;penam;penem | Cephalosporin | Cephamycin | fluoroquinolone |
| **Fragments #** | 11864 | | 24957 | 945 | 1262 | 675 |
| **ARGs #** | 723 | | 196 | 206 | 161 | 119 |
| **Class-specific features #** | 5 | | 62 | 1 | 1 | 1 |
| **most frequent class-specific features** | AAGC, GCAA | | CACTTTTAA, TCTGCTATGT, GATG, GTAT | TGGT | TACT | GGTG |
| **Class-specific features found in functional domains** | AAGC, GCAA | TGGA | GATG, GTAT, GGTGA, GCTGA, CACTTTTAA and TCTGCTATGT | NA | NA | GGTG |
| **Name of the functional domain** | β-lactamase class-A active site (PS00146) | β-lactamases class-B signature-2 (PS00744) | β-lactamase class-A active site (PS00146) | NA | NA | 1. Major facilitator superfamily (MFS) profile (PS50850)<br>2. ABC transporter integral membrane type-1 fused domain profile (PS50929)<br>3. ATP-binding cassette, ABC transporter-type domain profile (PS50893) |
| **ARGs containing the functional domain #** | 240 | | 195 | NA | NA | 14 |

#### Pseudocode for the MACI framework

```
Data:  fragment ←length (100)
if length > 100 then
slidingwindow(fragment)
end
Result: output →Class Number
Procedure:
def slidingwindow(fragment) :
  lengthfragments = [ ]
  for i in range(0, length(fragment) − 100) :
     lengthfragments.append(i : i + 100)
  return lengthfragments;
encoding ←fragment based encoding;
A ←1;
G ←0.75;
C ←0.25;
T ←0.5;
for class in classname:
  if length(class) ≥ 30000:
    modelclass1.append(class)
else:
    modelclass2.append(class)
classnames1=LabelEncoding(modelclass1);
classnames2 = LabelEncoding(modelclass2);
modelclass1 = [ ];
modelclass2 = [ ];
inputdata = [ ];
outputdata1 = [ ];
outputdata2 = [ ];
for fragment in encoding:
   classarray1 = np.zeros[size = (22)];
   classarray1 = np.zeros[size = (94)];
   if class in modelclass1:
       classarray1[classnames1] = 1;
```

```
   else:
       classarray2[classnames2] = 1;
   inputdata.append(fragment);
   outputdata1.append(classarray1);
   outputdata2.append(classarray2);
model1.layer1 ←100 neurons, activation = 'relu' type= 'Dense';
model1.layer2←400 neurons, activation = 'relu' type= 'Dense';
model1.layer3←200 neurons, activation = 'relu' type= 'Dense';
model1.layer4←22 neurons, activation = 'relu' type= 'Softmax';
model1.compile(loss = 'categorical crossentropy', optimizer= 'Adam', metrics=['accuracy'])
model1.train(input = inputdata, output = outputdata1, iterartions = 70)

model2.layer1 ←100 neurons, activation = 'relu' type= 'Dense';
model2.layer2←400 neurons, activation = 'relu' type= 'Dense';
model2.layer3←200 neurons, activation = 'relu' type= 'Dense';
model2.layer4←96 neurons, activation = 'relu' type= 'Softmax';
model2.compile(loss = 'categorical crossentropy', optimizer= 'Adam', metrics=['accuracy'])
model2.train(input = inputdata, output = outputdata2, iterartions = 70)

def classprediction(model1, model2, fragment) :
  output1 = model1.predict(fragment)
  output2 = model2.predict(fragment)
  max_value1 = max(output1)
  max_value2 = max(output2)
  if max_value1 > max_value2:
      index = find_index(output1, max_value1)
      output = LabelDecode(classnames1, index)
  else:
      index = find_index(output2, max_value2)
      output = LabelDecode(classnames2, index)
  return output
```
